# CHETAH: a selective, hierarchical cell type identification method for single-cell RNA sequencing

**DOI:** 10.1101/558908

**Authors:** Jurrian K. de Kanter, Philip Lijnzaad, Tito Candelli, Thanasis Margaritis, Frank C.P. Holstege

## Abstract

Cell type identification is essential for single-cell RNA sequencing (scRNA-seq) studies that are currently transforming the life sciences. CHETAH (CHaracterization of cEll Types Aided by Hierarchical clustering) is an accurate cell type identification algorithm that is rapid and selective, including the possibility of intermediate or unassigned categories. Evidence for assignment is based on a classification tree of previously available scRNA-seq reference data and includes a confidence score based on the variance in gene expression per cell type. For cell types represented in the reference data, CHETAH’s accuracy is as good as existing methods. Its specificity is superior when cells of an unknown type are encountered, such as malignant cells in tumor samples which it pinpoints as intermediate or unassigned. Although designed for tumor samples in particular, the use of unassigned and intermediate types is also valuable in other exploratory studies. This is exemplified in pancreas datasets where CHETAH highlights cell populations not well represented in the reference dataset, including cells with profiles that lie on a continuum between that of acinar and ductal cell types. Having the possibility of unassigned and intermediate cell types is pivotal for preventing misclassification and can yield important biological information for previously unexplored tissues.

## INTRODUCTION

Single-cell RNA-sequencing (scRNA-seq) is transforming our ability to study heterogeneous cell populations^1–6^. While tools to help interpret scRNA-seq data are developing rapidly^7–15^, challenges in data analysis remain^16^, with cell type identification a prominent example. Accurate cell type identification is a prerequisite for any study of heterogeneous cell populations, both when the focus is on subsets of a particular cell type of interest or when investigating the population structure as a whole^17–21^. The introduction of single cell RNA sequencing has paved the way for rapidly discovering previously uncharacterized cell types^22–24^ and this application too would greatly benefit from efficient identification of known cell types prior to focusing on new types.

Research into tumor composition presents an even more challenging setting, as the RNA expression profile of malignant cells is often different from any known cell type, as well as unique to the patient or even to the biopsy^25,26^. Malignant cells can sometimes be identified in scRNA-seq data^27^ but this is not always feasible or even possible, for instance with tumors that do not harbor easily identified copy number variations. In both cases, a first sign of the malignancy of cells in the sample is their imperviousness to classification, simply because their expression profiles do not resemble that of any known, healthy cell type.

Cell type identification in scRNA-seq studies is currently often done manually, starting by identifying transcriptionally similar cells using clustering. This is frequently followed by differential expression analysis of the resulting cell clusters combined with visual marker gene inspection^4,25,26,28–30^. Such manual cell type identification is time-consuming and often subjective due to the choice of clustering method and parameters for example, or to the lack of consensus regarding which marker gene to use for each cell type. Such analyses are becoming more complex given the fast-expanding catalogue of defined cell types^16^. Canonical cell surface markers are also not always suitable in scRNA-seq studies because the transcripts of these genes may not be measurable in the corresponding cell type owing to low expression or to degradation of the mRNA. This is aggravated by technical difficulties (drop-out) and, more generally, by the poor correlation between protein expression and mRNA abundances^23^.

Recently, a number of cell type identification algorithms have emerged to address these problems. Automated methods such as scmap^31^ and SingleR^32^ base their cell type call on comparisons with annotated reference data using automatically chosen genes that optimally discriminate between cell types. A good cell type identification method should be both sensitive and selective. That is, it should correctly identify as many cells as possible, while not classifying cells when based on insufficient evidence. If the cell being identified is of a type that is not represented in the reference, such misclassification can easily occur. This is a concern when studying malignant cells which are often too heterogeneous to include in the reference data. To avoid overclassification, methods such as scmap^31^ therefore leave cells unclassified if they are too dissimilar to any reference data.

Both the complete lack of classification as well as overclassification is unsatisfactory. For example, if the evidence for a very specific cell type assignment such as *effector CD8 T-cell* is not strong enough, a more general, less specific assignment such as *T-cell* may still legitimately be made and might still be useful. The reason for such an intermediate cell type assignment could be that the correct T-cell subtype is not part of the reference dataset, or even that there is insufficient read-depth for the more specific call to be made. An even more more interesting case is that of cells that are biologically of an intermediate type such as differentiating cells or cells undergoing transdifferentiation.

Here we present CHETAH (CHaracterization of cEll Types Aided by Hierarchical clustering), an algorithm that explicitly allows the assigningment of cells to an intermediate or unassigned type. The unassigned and intermediate types are inferred using a tree that is constructed from the reference data and which guides the classification. CHETAH’s classification is a stepwise process that traverses the tree and, depending on the available evidence, ends at one of the reference cell types or halts at the unassigned or one of the intermediate types. CHETAH is able to correctly classify published datasets and, in comparison to other methods it performs equally or better when considering cells whose type is represented in the reference data. For cells of an unknown type, CHETAH is more selective, yielding a classification that is as fine-grained as is justified by the available data. The benefit of calling unassigned and intermediate types is highlighted in several tumor datasets, showing CHETAH is consistently selective. This makes CHETAH a powerful tool for identifying cells that are not in the reference, such as malignant tumor cells, novel or intermediate cell types. The latter is shown in an analysis of published pancreas datasets, where a manifest expression gradient of cells with types varying between *acinar* and *duct cell* is described. CHETAH is implemented in R^33^ and is available at github.com/jdekanter/CHETAH. It comes with an extensive Shiny^34^ application that makes exploration of the cell type identification process and the gene expression differences that support the classification very intuitive. CHETAH has been created bearing tumor analyses in mind, but as is demonstrated, it also complements existing methods for exploring previously uncharacterized non-cancerous tissues and cell types.

## METHODS

An outline of the CHETAH algorithm is depicted in Figure 1. First a hierarchical classification tree is constructed from the reference scRNA-seq data. This tree guides the classification in a way that resembles the use of a taxonomic key. At each step of the process the cell to be classified is correlated to the expression profiles of the reference data. Each correlation is compared to the distribution of correlations of the reference cells per cell type to assess whether there is enough evidence to allow this cell to take the next step. If not, further classification of the cell stops and it is marked as *unassigned* if the evidence runs out at the top of the tree, or as intermediate if this happens within the classification tree. Classification also stops when a cell reaches one of the leaves of the tree.

**Figure 1.**
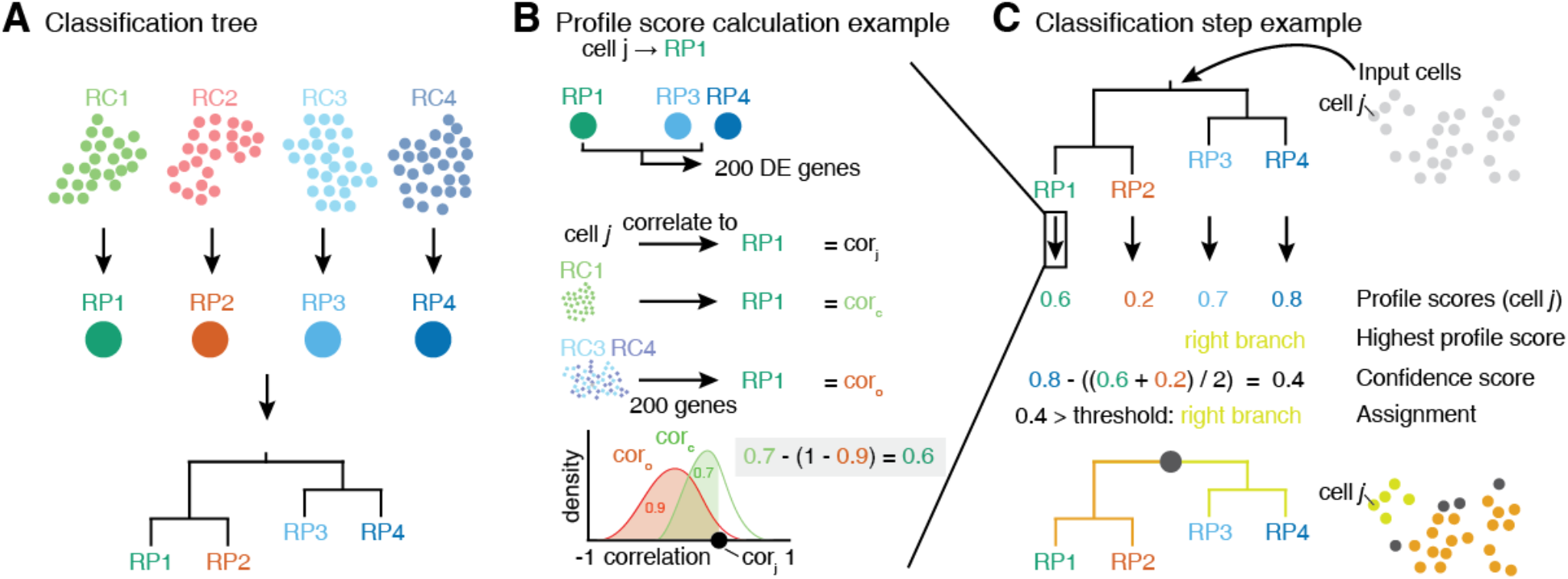
The CHETAH algorithm. For details see Methods. (A) Reference profiles (RPs) are created for each cell type by averaging the expression profiles of all reference cells (RCs) per type. A classification tree is computed from these RPs using Spearman correlation and average linkage. The input cells are classified in a top-to-bottom, stepwise process traversing the tree. In each node, the input cells are assigned to the left or right branch, or to the node itself, depending on the input cell’s profile and confidence scores (B) For each input cell *j*, a profile score expressing cell *j*’s similarity to each candidate RP is calculated (shown for RP1). To this end the fold-changes in expression of all genes between the candidate RP (here: RP1) and the mean of all the RPs in the other branch (here RP3 and RP4) are calculated. The 200 genes with the highest absolute fold-change are selected and used to correlate each individual reference cell to RP1. This results in a distribution of expected correlations of cells of the current candidate cell type. Likewise, the distribution of correlations of the union of all cell types under the other branch (here RC3 and RC4) to RP1 is calculated. The profile score of input cell *j* is based on the cumulative probabilities of cell *j*’s correlation within these two distributions. (C) Lastly, cell *j* is assigned to the branch containing the RP with the maximal profile score (here RP3+4) but only if the confidence score for this assignment is large enough. If not, the cell’s type is *unassigned* (if it is the top node of the tree) or one of the *intermediate types* (if it is further down). The confidence score is based on all profile scores under the current node.

### Reference data

In order to classify input cells CHETAH requires scRNA-seq reference data along with cell type labels for each reference cell. The reference data needs to be normalized to an identical total number of transcripts per cell and should be expressed in log-scale. Malignant cells are best left out of the reference because they are too ill-defined and patient-specific^35^. In all the reference datasets used here, such cells cluster by patient whereas non-malignant cells largely cluster by cell type. The reference must contain at least 10 cells per cell type to adequately represent its transcriptional program as well as its variance. The reference must contain at least 10 cells per type to adequately represent its transcriptional program as well as its variance. Between 10 and 100 is optimal. More than 100 cells per cell type is superfluous and increases the computational burden.

Unless stated otherwise, the reference dataset used here consists of a combination of datasets of colorectal^36^, breast^37^, melanoma^25^ and head-neck tumors^26^. The data for all these studies was generated using the Smart-Seq2 platform. The cell types of the reference dataset were based on published manual classification of cell clusters using marker gene expression. The melanoma and head-neck studies discuss the T-cells in terms of their *CD4*+, *CD8*+ and *T-reg* subtypes but not all of these labels are available for all the cells in the online material of these publications. These reference cells were therefore classified manually using the same marker genes as used in the publications. Cells of a dataset being classified are of course excluded from the reference. When comparing CHETAH and SingleR^32^ results, the latter was run with averaged single-cell data because SingleR uses bulk, rather than single-cell expression profiles as its reference.

### Classification tree

The first step is to create a reference profile (RP) for each cell type in the set of reference cells by averaging, for each cell type, the logged gene expression over all cells of that type (Figure 1A). The RPs are subsequently clustered hierarchically using Spearman correlation and average linkage to obtain the classification tree.

### Hierarchical classification

The classification of input cells proceeds in a stepwise fashion, from the root to the leaves of the classification tree. At each step, the branch is selected that contains the reference cell type most likely to be the correct one, but the classification stops if the confidence in this decision becomes too low (see confidence score below). As described under profile score, the choice of the most likely cell type and therefore which branch to choose, is based on the cell’s similarities to each of the individual RPs under each branch. The similarity of a cell to a RP under consideration (called the candidate RP), in the branch under consideration (called the candidate branch), is always in relation to all the RPs under the (so-called) other branch. During the classification process, only the leaf node data (i.e. from all cells of a particular reference cell type) are used. Any details of the tree topology under either branch are ignored, i.e. no hypothetical expression profiles are inferred for the intermediate tree nodes. After calculating the cell’s similarities to all RPs under both branches, the cell is assigned to the branch that contains the cell type to which it is most similar, provided the evidence is strong enough based on the confidence score.

### Feature selection

The similarity of a cell to a reference profile is based on their Spearman correlation. This correlation is calculated using the subset of genes that best discriminates between the *candidate* RP in the *candidate branch,* and the average expression profile of the *other branch* as a whole. (The latter is calculated as the mean of all RPs under that branch). The selection of the best subset of genes, a process known as feature selection, is not critical and good results are achieved when simply using the 200 genes that have the largest absolute fold-change between the *candidate RP* and the average expression profile of the *other branch*. It is important to note that the feature set, i.e. the subset of genes used to calculate similarities, is generally different for each RP and for each node of the classification tree.

### Similarities

The similarity of a cell to a RP in the candidate branch is of course reflected in their correlation, but the values of these correlations to the various RPs cannot be directly used for comparisons. The reason is that the subset of genes used for each correlation is generally different for each RP and for each node. The similarity of an input cell *j* to candidate RP *x* is therefore cast in relative terms by using the *cumulative probabilities* of this correlation within two different *distributions* of correlations. The first one is the distribution of self-correlations, that is, the distribution of the correlations of the individual cells constituting the *candidate* RP to that *candidate* RP itself. These self-correlations represent the typical correlation values for a cell that is really of that type. The second distribution is that of the nonself-correlations. They are the correlations, again to the *candidate* RP, of all the individual reference cells under the *other* branch. They represent the correlation values that can be expected for cells that are *not* of any type under the candidate branch. By contrasting the two cumulative probabilities a profile score is obtained that robustly points the way through the classification tree.

### Profile scores

The two cumulative probabilities just defined are used to define the profile score *P*_*x*_(*j*), representing cell *j*’s similarity to *candidate* RP *x*, as follows:

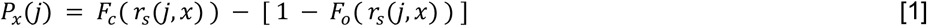

with

*r*_*s*_(*j, x*) the Spearman correlation of input cell *j*’s expression with candidate reference profile *x*
*F*_*c*_(*r*_*s*_(*j, x*)) the cumulative probability of *j*’s correlation within the distribution of self-correlations *r*_*s*_(*k, x*), that is, of all reference cells *k* of type *x* with their ‘own’ candidate reference profile *x*
*F*_*o*_(*r*_*s*_(*j, x*)) the cumulative probability of *j*’s correlation within the distribution of correlations *r*_*s*_(*l, x*) of all reference cells *l* under the *other* branch, again with reference profile *x*

The profile score *P*_*x*_(*j*)has a value between 1 and −1 and is, in a particular node, a measure for the likelihood that cell *j* is of type *x*. A value of 1 means that cell *j* is much more likely to be of type *x* (and therefore belong to its branch) rather than any of the types in the other branch and, conversely, −1 represents the lowest likelihood of this being so, and therefore cell *j* is much more likely to belong in the other branch. In each node, one set of genes is selected for each RP under that node. This gene set is used for all the correlations (of both input and reference cells) needed to calculate the profile scores. Note that due to the different gene subsets used in each step of the tree traversal, the most similar RP for a cell may change during the steps of the classification process. E.g., during the first few steps a cell that in reality is of type *CD4 T-cell* could initially and, incorrectly, appear more similar to a *CD8 T-cell* than to the expected *CD4 T-cell* type. This would however still lead to the correct branch choice, namely that of all *T-cells*. In later steps the similarity to the actual *CD4 T-cell* type would become strongest, guiding the cell to a correct final *CD4 T-cell* label.

### Confidence score

Each input cell is assigned to the branch containing the candidate reference cell type for which it has the highest profile score. This assignment represents the choice between the left and right branch, but a key design goal of the algorithm is its ability to stop classification at an intermediate node. The choice for each cell *j*, between stopping classification or continuing to the next round, is based on its confidence score *C*(*j*)defined as

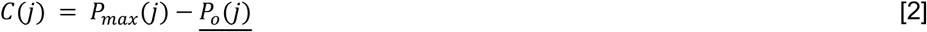

with *P*_*max*_(*j*) the highest profile score for cell *j* in the branch about to be chosen and 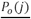 the mean of the profile scores in the *other* branch, i.e. the branch *not* containing the reference profile having the highest profile score (Figure 1C). Expression [2] is always positive because branches leading to a negative score are by definition never chosen by the algorithm. The confidence score is a measure for the evidence to assign a cell to a branch, with 2 representing maximal evidence, and 0 representing no evidence. The confidence score has an easy explanation. If it is close to 0, the best candidate cell type in the branch about to be chosen is as good as the average of the cell types in the other branch. This implies that there is no basis to justify the choice between *either* branch, so *none* should be taken and classification of the cell should therefore stop in the current node. In contrast, a large score represents good support to continue the classification because there is a cell type in the candidate branch that has a much better profile score than the average profile score of the other branch. By default, cells are assigned to the branch if the confidence score is higher than 0.1, but different values can also be specified in the algorithm’s parameters. Cells that remain in a non-leaf node of the tree are called *unassigned* or of *intermediate type* whereas cells assigned to a leaf-node are of a *final type*. The labels for the *intermediate types* are generated automatically (*Node1, Node2*, etc.) but biologically meaningful names such as *T cell* can often readily be given. By choosing a cut-off greater than 0.1, only the more confident calls will be made, hence more cells will be labeled as being *unassigned* or of *intermediate type*. Conversely, by lowering the confidence cut-off, the algorithm will classify more cells to a *final type*, however such calls are supported by less evidence. A cut-off of 0.0 forces the method to classify *all* cells to a *final type*, as is exemplified later. The above stepwise calculations of the profile scores and confidence scores yield an elegant and, importantly, transparent algorithm.

## RESULTS

The CHETAH algorithm can be recapitulated as follows. Reference cell types are hierarchically clustered into a classification tree which functions as a taxonomic key for the cell type identification process. The cells to be classified are shunted from the root of this tree to its leaves, but only to the most specific tree node that is still supported by the available evidence, as quantified by a robust measure of confidence. Cells for which this confidence runs out are typically of a type that is not present in the reference, and are said to be either *unassigned* or of an *intermediate type*. Note that there are several intermediate types, each corresponding to one of the internal nodes of the classification tree.

CHETAH’s accuracy is investigated by comparing its classifications with published cell type labels. The aim is to reproduce these using only the reference data. Since the accuracy might be lower if the scRNA-seq technology of the input data and the reference differs, cross-platform results are also examined. CHETAH is subsequently compared to other cell type identification methods and the effectiveness of the intermediate cell type assignments is also demonstrated in an analysis of previously published pancreas datasets.

### Accuracy

The performance of CHETAH is first evaluated by applying it to melanoma^25^ and head-neck^26^ cancer datasets. The classifications of the these datasets is shown in Figures 2A,B (Melanoma), S1A,B (Head-Neck) and summarized in Table 1. Since the reference datasets do not contain malignant tumor cells, such cells should not be classified to any *final type,* but as *unassigned* or any of the *intermediate types* instead. CHETAH correctly classifies practically all (mean > 99%) malignant cells as *unassigned* or *intermediate* types. Note that in the published data the classifications were manual while the identification of tumor cells was based on estimated copy number variations. In contrast, CHETAH’s type assignments are fully automatic and the aberrant nature of the malignant cells is indicated by their classification as *unassigned* or *intermediate*. This selectivity is an important quality of the method, essential for preventing the type of misclassification that readily occurs when methods forcefully assign every cell to a type regardless of the evidence. Selectivity is especially relevant when dealing with tumor samples, as well as with samples containing cell types not present in the reference dataset.

**Table 1.** Percentages of cell type labels as inferred by CHETAH, compared with the published cell types. The column *correctly unassigned* shows the percentage of cells of a type that was absent from the reference that were classified as *unassigned* or any of the *intermediate type*s. The other columns refer to sample cells of a type represented in the reference that should therefore be assigned and contain percentages of cells of final or intermediate type, summing to 100%. The term *lineage* refers to the classification tree determined by CHETAH.

| percentage dataset | correctly unassigned | identical final type | intermediate type, same lineage | intermediate type, different lineage | incorrectly unassigned | different final type |
| --- | --- | --- | --- | --- | --- | --- |
| Melanoma<sup>25</sup> | 99.5 | 86.9 | 4.1 | 0.8 | 0.5 | 7.7 |
| Head-Neck<sup>26</sup> | 89.2 | 71.2 | 14.6 | 1.1 | 1.9 | 11.2 |
| Ovarian<sup>19</sup> | 99.3 | 79.9 | 9.0 | 0.7 | 6.8 | 3.6 |

**Figure 2.**
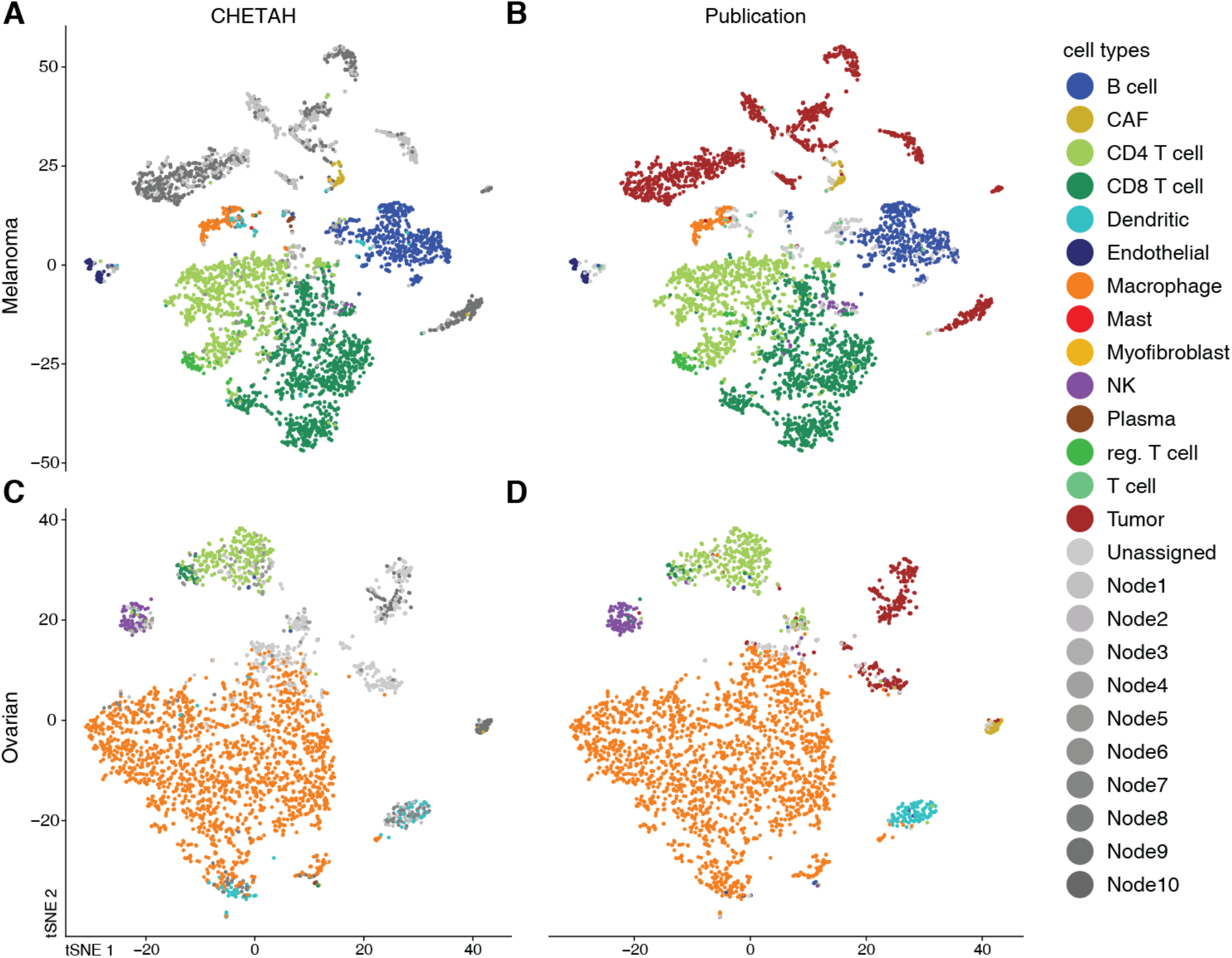
CHETAH’s classification of two tumor sample datasets is nearly identical to the published manual classification. The t-SNE plots depict each cell as a dot, with the colours representing the inferred cell type shown in the legend. Gray colours indicate intermediate cell types which are labeled automatically as *Node1, Node2,* etc. For the corresponding classification trees see Figure S2. Rows of panels: datasets classified (Melanoma: Tirosh *et al.*^25^; Ovarian: Schelker *et al*.^19^); columns: classification method.

CHETAH classifies the majority (mean 79%) of non-malignant cells the same as in the original publication. Of the cells classified differently, the majority (mean 61%) are classified as an *intermediate* type. In the inferred classification tree these *intermediate* assignments are overwhelmingly in the correct classification lineage (85%, 91% and 95% for Melanoma, Head-Neck and Ovarian respectively). Only a small number of cells are labeled differently by CHETAH. For many of them there is in fact strong evidence from established marker gene expression that the assignment from CHETAH is correct (Figures S3, S4). Taken together, these results show that the selective approach works well. Cells of an established type that are present in the reference dataset are classified correctly. Samples cells of a new or aberrant type, not represented in the reference dataset are either not assigned to a type or are classified as an intermediate type, an outcome that should indeed be regarded as a pointer for a more detailed inspection.

### Cross-platform classification

The data from the Melanoma and Head-Neck studies were obtained using Smart-seq2^38^ and were also classified using reference data originating from the same platform. To evaluate CHETAH’s performance across platforms, an ovarian tumor dataset^19^ produced on the inDrops platform^39^ was analyzed with CHETAH using the Smart-Seq2-based reference. The results, presented in Figure 2C,D and Table 1, show a performance similar to that obtained within one platform. This robustness is probably due to the use of rank-based similarities, implying that other combinations of scRNA-seq technologies will likely yield similar good results.

### Comparison with existing methods

The important challenge of cell type identification has recently also started to be addressed through the development of other automated approaches. CHETAH was therefore next compared to the state-of-the-art methods CaSTLe^40^, scmap^31^ (both versions, i.e. scmap_cell and scmap_cluster) and SingleR^32^ by running these programs with standard settings on the Ovarian, Melanoma and Head-Neck datasets (Figure 3A). To evaluate the performance also on non-tumor tissues, two pancreas datasets, Pancreas1^17^ and Pancreas2^41^ were included and mutually classified using the other as the reference. To avoid bias the same reference data was used for all classification methods. The ground truth for the classifications are the cell type labels from the original publications, but without the malignant cells from the tumor datasets. They are not part of the reference data and should therefore be considered as an unknown cell type and remain *unassigned* or *intermediate*.

**Figure 3.**
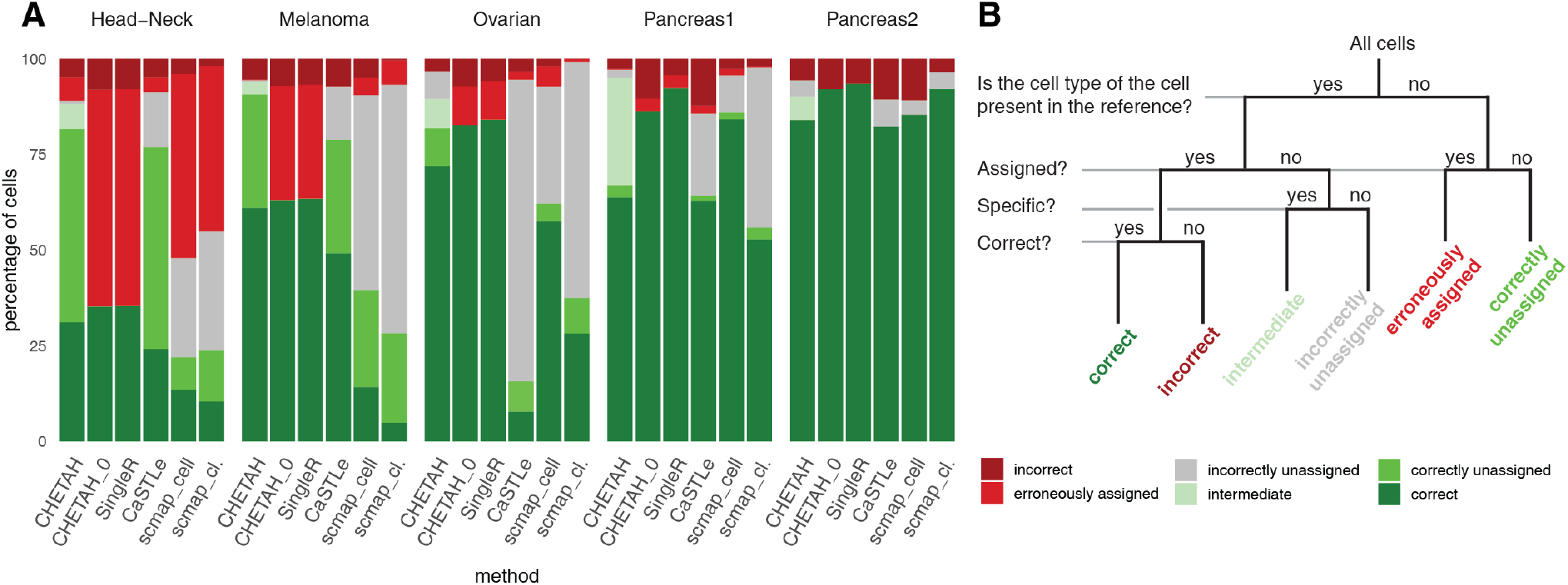
CHETAH compared with other methods (bottom labels), across five datasets (top labels). For scmap, both the ‘cell’ mode (scmap_cell) and ‘cluster’ mode (scmap_cl.) where evaluated. CHETAH was run with default settings, but also with a zero confidence score threshold (CHETAH_0), thus forcing it to classify all cells to a final type. (A) Percentages of cells per classification result category as shown in (B). (B) Classification result categories used in (A).

To compare methods, two classes of input cells can be distinguished, namely (1) the cells that are of a type that is present in the reference and (2) cells for which no reference is available (Figure 3B). For the first class it is meaningful to assess thecorrectness of the classification, because an optimal method should correctly identify all such cells. Those cell type inferences can therefore be *correct* or *incorrect,* corresponding to the true and false positives respectively (Figure 3B). In addition, the categories *intermediate* or *unassigned* are allowed, to accommodate methods such as scmap and CHETAH that produce *intermediate* and *unassigned* calls. The second class of input cells, those of a type absent from the reference, should *not* be classified by an optimal cell type identification. These are therefore divided into *correctly unassigned* cells which can be considered true negatives, and their false positive counterparts, here called *erroneously assigned*, i.e. cells that were, but should not have been, classified.

In the cancer datasets, CHETAH generally outperforms other methods (Figure 3A) in terms of combined true positives (correct assignments) and true negatives (cells correctly left unclassified). This is particularly important for studies of cancer since malignant cells are typically very patient specific and would almost always be misclassified by greedy methods. SingleR, having no classification cut-off, always classifies *all* cells to a final type, leading to a large number of *erroneously assigned* cells in cancer samples with many malignant cells (Figure 3). For example, both the cancer-associated fibroblasts (CAFs) and malignant cells are all classified as CAFs by SingleR. In datasets containing many unknown cells such as the malignant cells in the cancer samples, such approaches would therefore require very careful post hoc inspection of the classification on a per cell or per cluster basis, an approach that automated methods are meant to obviate. The selective nature of CHETAH makes the analysis much more efficient. As anticipated, forcing CHETAH to become greedy and classify all cells by applying a confidence score threshold of 0, yields a performance almost identical to SingleR’s (Figure 3).

In contrast to the cancer datasets, the pancreas data are less complex, containing cell types with strong differential gene expression and few unknown cells. Note that a perfect classifier should leave none of the cells in the Pancreas2 dataset unidentified, because all its cell types are represented in the Pancreas1 reference. The converse is not true because for some of the cell types no distinction is made in Pancreas1. This is the reason that all the methods perform better on Pancreas2 (Figure 3A). In this comparison on non-cancer datasets, CHETAH’s forte of rarely classifying cells without sufficient resemblance to the reference cell types is diminished. This results in a performance similar to that of the other methods (Figure 3). However, as is exemplified below, the inclusion of an intermediate assignment can have benefits for such datasets too.

### Intermediate types

In data from tumor samples the classification to an *intermediate type* suggests, by exclusion, that a cell is aberrant and therefore potentially malignant. The position in the classification tree, of the node of an intermediate type may shed further light on the biology of these cells. For example, in the Melanoma and Head-Neck datasets, 54% and 74% respectively of the malignant cells, classify to the node directly above *endothelial*. This suggests that the expression pattern of these cells shares characteristics with endothelial and fibroblast types (see Figures S2A,B for the classification trees). Conversely, these cells display no affinity with the hematopoietic lineage, which is consistent with these tumors not being of hematopoietic origin. Classification to an intermediate type in combination with the position in the classification tree is therefore useful for analysis of cancer datasets.

Assignment to an intermediate cell type can also be useful in non-cancer datasets. This is demonstrated by two examples. In the Pancreas1 dataset, two kinds of stellate cells were originally identified, both of which are of mesenchymal origin^42^. *PDGFRA* and *RGS5* were applied as marker genes for *activated* and *quiescent stellate* cells respectively. Pancreas2 only contains the more general label *mesenchymal*, and the corresponding cells only exhibit expression of *PDGFRA* but not *RGS5* (Figure S5), implying that these reference cells more closely resemble *activated* rather than *quiescent stellate* cells. When CHETAH classifies the Pancreas1 dataset using the more limited Pancreas2 reference data, it correctly identifies the Pancreas1 *activated stellate* cells as *mesenchymal* while leaving the *quiescent stellate* cells *unassigned*, or assigning them to the node directly above the *endothelial* and *mesenchymal* types (Figure S7B), correctly determining that these cells are of a mesenchymal type not represented in the reference.

### Acinar - duct cell gradient in pancreas data

Another useful consequence of allowing an intermediate type is exemplified in Figure 4. Some cells in a cluster identified as *acinar* in the Pancreas2 publication are labeled *ductal* by CHETAH (Figure 4A), while conversely the cluster called *ductal* in the Pancreas1 study is partly classified as *acinar* (Figure S7A). The presence of these mixed acinar-ductal groups in both datasets suggests a shared underlying phenomenon. *Acinar* and *ductal cells* arise from the same progenitors and are closely related^43^. They are separated by only one node in CHETAH’s classification tree (Figures 4B and S7B), which is the intermediate type to which CHETAH assigns the remaining cells of these clusters. When visualizing the profile score for *duct cell* in this intermediate node (arrows in Figures 4B and S7B), a smooth gradient is clear in both clusters (Figures 4C and S7C).

**Figure 4.**
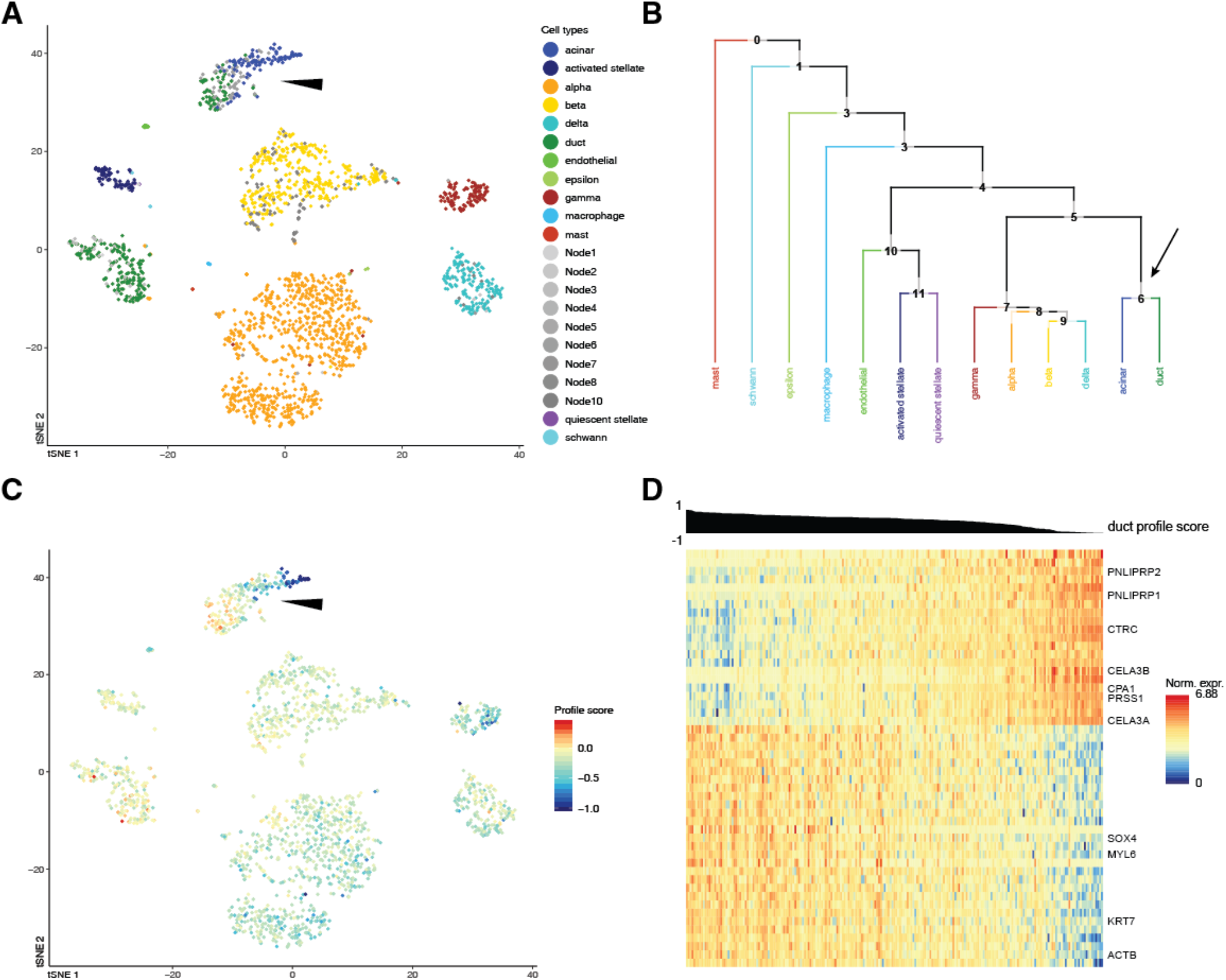
CHETAH identifies opposing gradients of duct and acinar cell marker genes in the Pancreas2 dataset^41^. (**A**) t-SNE plot of the Pancreas2 dataset as classified by CHETAH, with colours representing the inferred cell types. The arrowhead indicates a population that was labeled as *acinar cell* in the publication, but is classified to a mixture of *duct cell* (blue), *acinar cell* (green) and *intermediate Node 6* (grey) by CHETAH. (**B**) The classification tree used for A, based on the Pancreas1 dataset. The arrow indicates the acinar/ductal intermediate node (*Node 6*) for which the profile score of duct cells is shown in C. (**C**) As B, but with all cells coloured by the profile score for *ductal cell* in *Node 6*. The cells in the cluster of interest show a gradient of the profile score. (**D**) Heatmap showing the normalized expression of the genes (rows) used by CHETAH to calculate the profile score plotted in C, for the cells (columns) in the cluster indicated by the arrowhead in panel A. Only genes having an absolute correlation > 0.5 with the profile score are shown. The cells are sorted by the *duct cell* profile score in *Node 6* which is shown above the heatmap. Well-known acinar (top) and ductal marker genes (bottom) are labeled (see main text). For the heatmap with all genes annotated see Figure S6.

A heatmap of the expression of the genes most strongly (anti)correlating with this profile score shows the well-known cell type markers for these cell types (Figures 4D and S7D). These cell type-specific markers again exhibit a gradient of decreasing ductal and increasing acinar expression. Among the negatively correlating genes are acinar markers like *CPA1, PRSS1* and *CTRC*^44,44^ and among the positively correlating genes are pancreas duct cell markers like *KRT7* and *KRT19*^46^. A similar gradient in the expressions of genes having unusually large loadings in the first principal component of their ductal cell population has been reported previously^17^. This is a different manifestation of the fact that, for these cells, there is no dichotomy between *acinar* and *ductal*. Instead, the type of these cells is best described as lying on a continuum between *acinar* and *ductal*. The intermediate type assignment and profile score provide a direct and intuitive visualization highlighting such cases and the utility of the approach taken by CHETAH.

## DISCUSSION

Classification of cell types in scRNA-seq data is an essential step that was by necessity initially performed manually^27,25,26,19^. Owing to the subjective and time-consuming nature of manual approaches, automated approaches have recently been developed^31,40,32^. CHETAH has several features which work in its favour. Importantly, it compares input cell data with real, rather than imputed reference cell profiles. Moreover, besides using correlations, the classification decision is also based on a confidence score determined by the degree to which an input cell matches the expression variance embodied by the cumulative distribution function of the correlations to the reference cells. This facilitates the highly selective nature of CHETAH, underlying the ability to classify cells as specifically as the input and reference data allows, but without greedy over-classification, as controlled through the confidence score threshold. One consequence is the assignment to intermediate or unassigned cell types for input cells not present in the reference data. The assignment to an intermediate or unassigned type is essential to prevent overclassification and acts as an automated flag to more closely inspect such cells. The importance of this is evident both from the tumor datasets for which the method was initially devised, but also for non-cancer datasets as is also exemplified. In the tumor datasets analysed here practically all malignant cells were classified to intermediate types. Although genetic lesions such as copy number variations can be used to identify malignant cells^25^, this does represent an additional step. Moreover, such aberrations are not necessarily present (as in many pediatric tumors^43,44^) and/or may not be readily detectable. Automated highlighting of malignant tumor cells by CHETAH through classification as an intermediate or unassigned cell type is a significant improvement compared to blind misclassification. CHETAH’s confidence threshold can be adjusted to the needs of the dataset at hand, making it a flexible tool for research. CHETAH is made available as an R^33^ package, useful for application in conjunction with tools such as SCENIC^9^, Scater^10^, Census^11^, Monocle^12^, Seurat^13^, MERLoT^14^ and CellBIC^15^. The CHETAH package additionally includes a Shiny^34^ app for intuitive visualisation of the type labels, profile and confidence scores in a t-SNE^47^ plot, as well as the inferred classification tree and expression heatmaps of discriminatory genes.

In the pancreas datasets, CHETAH uncovers a group of cells exhibiting a gradient of profiles between *acinar* and *ductal*, previously suggested to be centroacinar cells^10^. An alternative explanation is that these are acinar cells undergoing acinar-to-ductal transdifferentiation or metaplasia (ADM)^34^. This is commonly seen in acinar cells that, like those in both pancreas studies discussed here, are cultured for several days^35^ or subjected to stresses or injury^36^. Subtle phenomena such as the *acinar-ductal* gradient are easily overlooked by greedy methods and especially by (manual) methods that assign the same cell type to all cells of one apparent cluster. Classification of cells from diverse tissues and diseases will become easier with the increasing availability of scRNA-seq datasets. Efforts like the Human Cell Atlas (HCA)^24^ are aimed at generating scRNA-seq datasets for almost each (healthy) tissue and cell type. CHETAH’s accurate handling of unknown cell types should prove useful in discovering novel cell types in such data. Conversely, the annotated HCA data would be very suitable as a reference for CHETAH.

CHETAH is not limited to the use of scRNA-seq and can likely be used with other quantitative single cell data such as those obtained using DNA accessibility^48,49^, chromatin state^50^, methylome^51^, epitope^52^ or RNA velocity^53^ sequencing methods, provided sufficiently rich reference data is available. Although the full range of single cell genome-wide approaches can be expected to increase further in the near future, the need for methods such as CHETAH that improve the ease and precision of the analysis of the resulting data is evident.

## SOFTWARE AVAILABILITY

CHETAH is available at github.com/jdekanter/CHETAH. All scripts that are needed to perform the analyses mentioned in this paper and to create the t-SNE plots using Seurat^13^ are deposited at github.com/jdekanter/CHETAH_paper_figures.

Supplementary Data are available at NAR online.

## ACKNOWLEDGEMENTS

We thank members of the Holstege, Kemmeren and Molenaar groups at the Máxima Center for insightful discussions and comments.

## Supplementary figures

**Figure S1.**
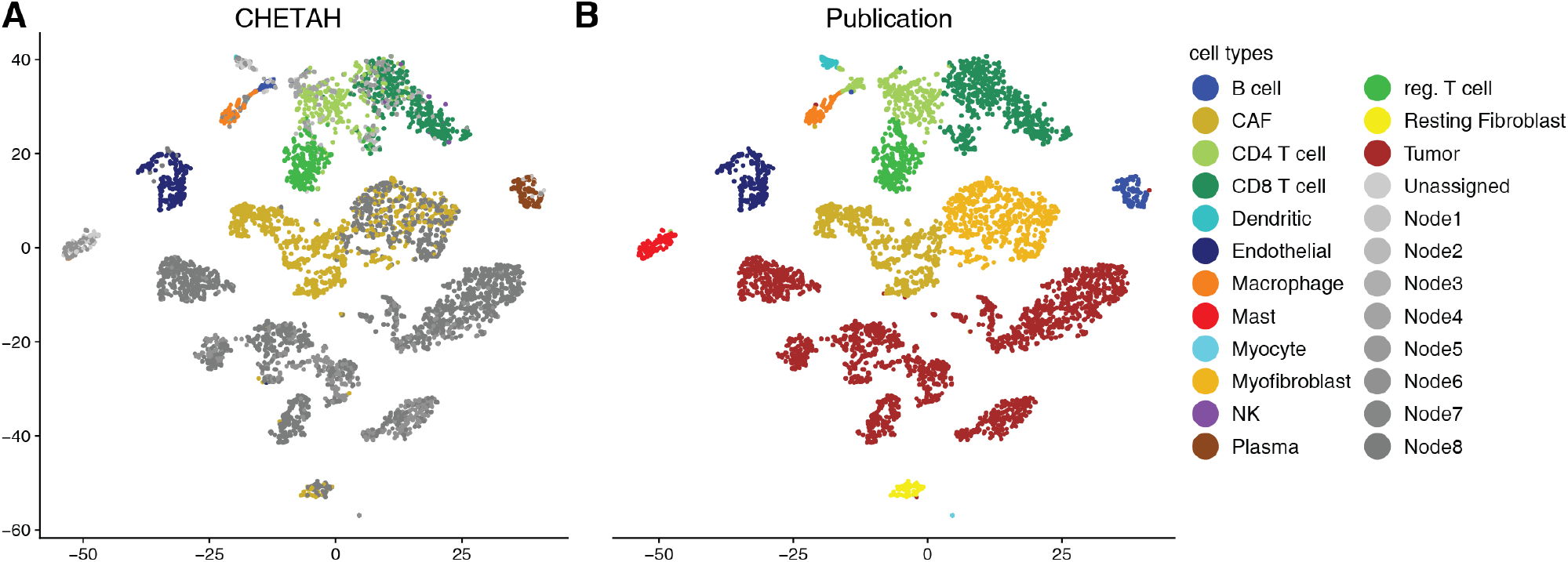
Classification of the Head-Neck tumor data (Head-Neck dataset: Puram *et al*., 2017, reference 18 in main text). Details as in main Figure 2.

**Figure S2.**
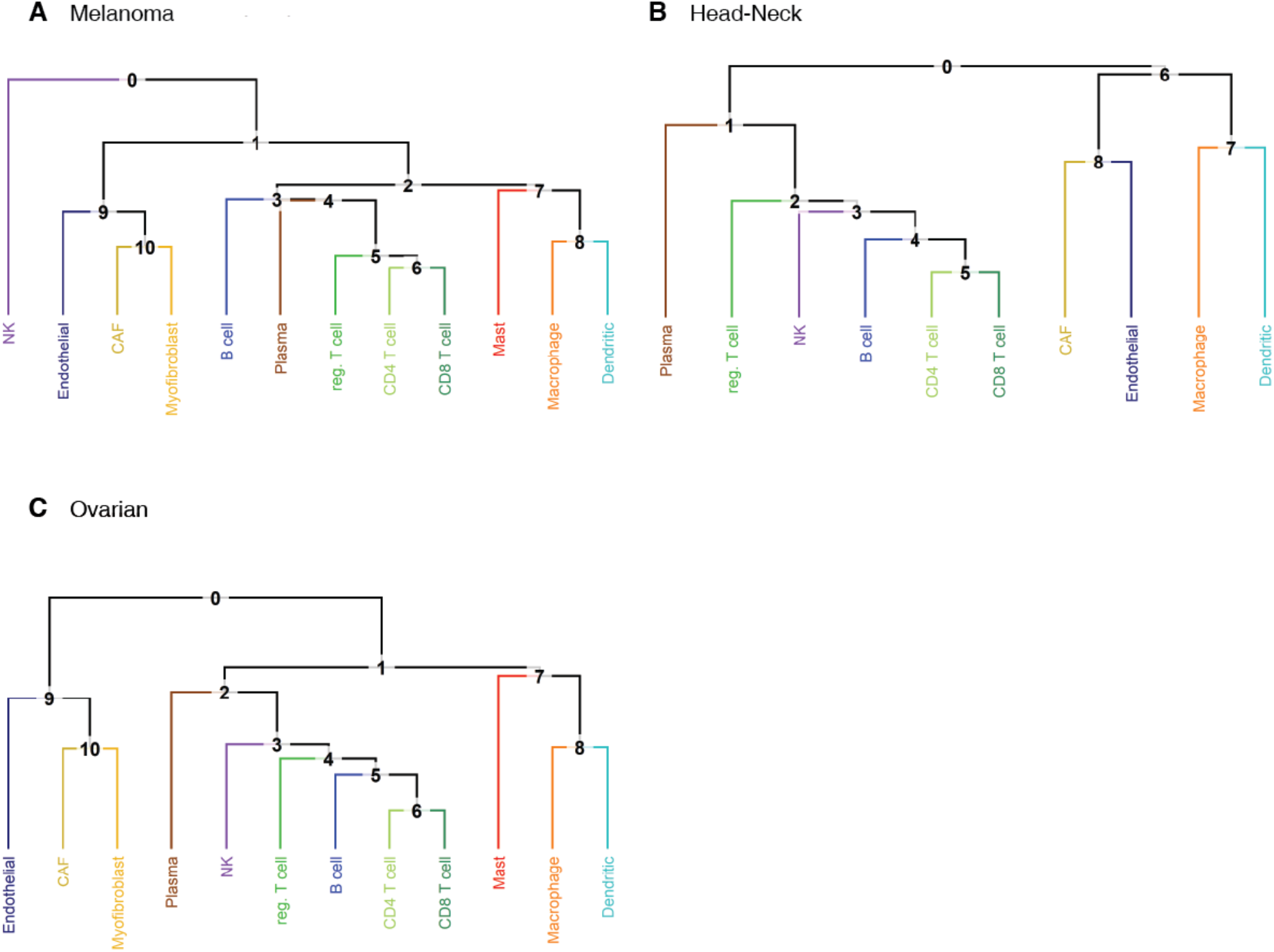
The classification trees used for the tumor datasets. Note that the trees are similar and reflect known biology. Small differences can arise due to a limited number of reference cells available (e.g. just 14 NK reference cells in the Melanoma classification tree). In the Melanoma and Head-Neck datasets, respectively 54% and 74% of the cells labeled as malignant cells classify to the node above *cancer associated fibroblasts* and *endothelial*.

**Figure S3.**
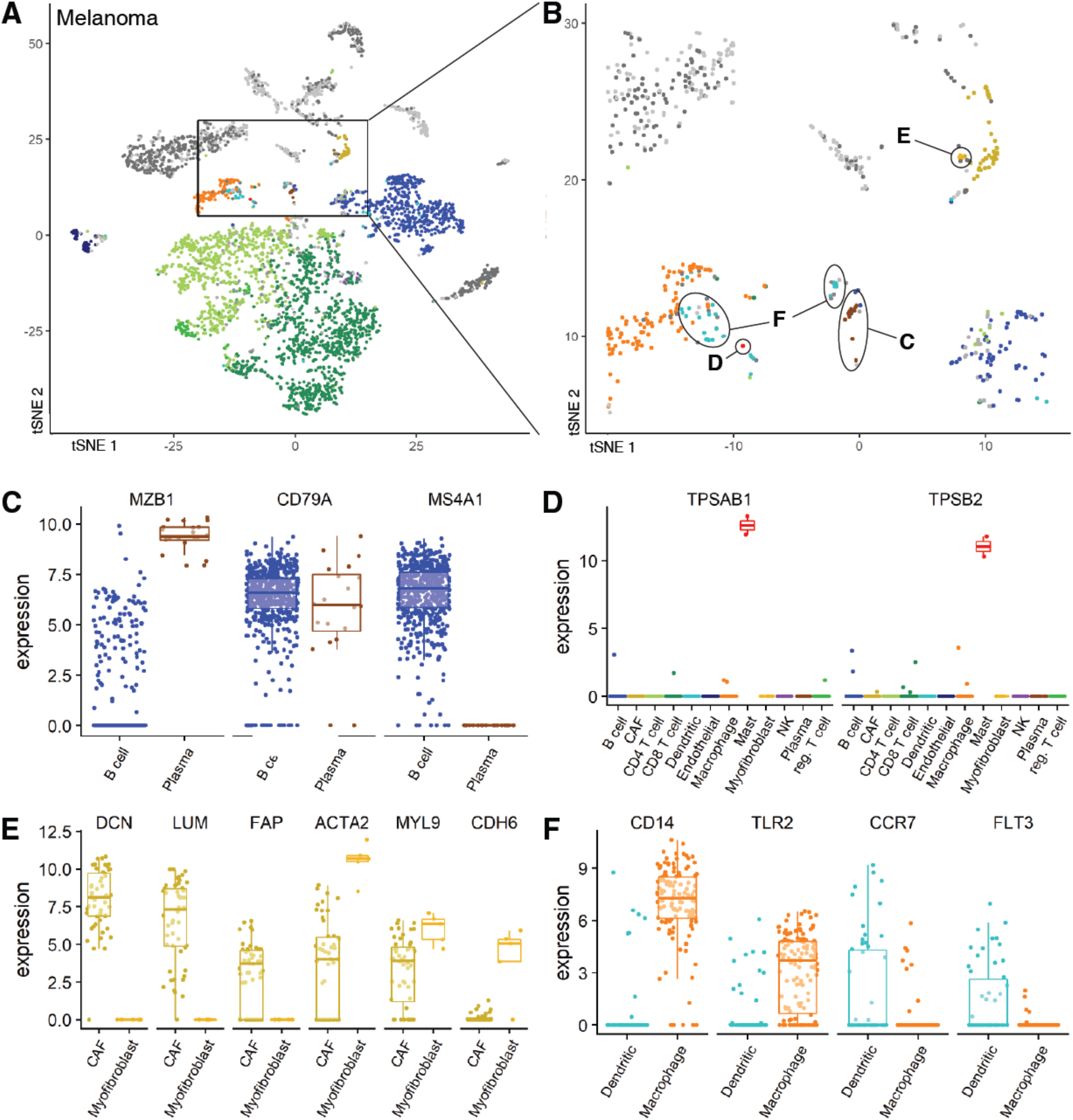
When CHETAH classification differs from the one assigned in the Melanoma study (Figure 2A), marker gene expression suggests CHETAH’s classification is more plausible. t-SNE plots of (**A**) the Melanoma dataset and (**B**) a zoom-in of this dataset. Details are as in main Figure 2. The arrows point to the discrepantly classified populations with letters C-F indicating the panel in which the marker gene expression of this population is plotted. (**C-F**): For each cluster shown in panel **B**: boxplots of marker gene expression (number of transcripts) of the marker genes (shown on top) for cells of the conflicting cell types (CHETAH classification shown at the bottom). colours are the same as in panels **A,B.** (**C**) Most of the cells previously classified as B cells in the Melanoma dataset are probably naive B cells, but some (encircled C in panel B) are more likely plasma cells. Most of these cells express B cell marker *CD79A*. Most plasma cells express activation marker *MZB1* but lack expression of *CD20/MS4A1* which goes down upon activation. The naive B cells express these two genes in opposite manner. (**D**) Two previously unclassified cells are probably mast cells. These are the only cells that highly express mast markers *TPSAB1* and *TPSB2*. (**E**) Previously unclassified cells in the cancer-associated fibroblasts (CAFs) cluster are probably myofibroblasts. These cells express none of the CAF markers (*DCN, LUM, FAP*), while they express higher levels of actin and myosin genes compared to the CAFs. (**F**) Previously unclassified cells are probably dendritic cells. Few of these cells express macrophage markers (*CD14, TLR2*), while they are enriched for dendritic markers (*CCR7, FLT3*).

**Figure S4.**
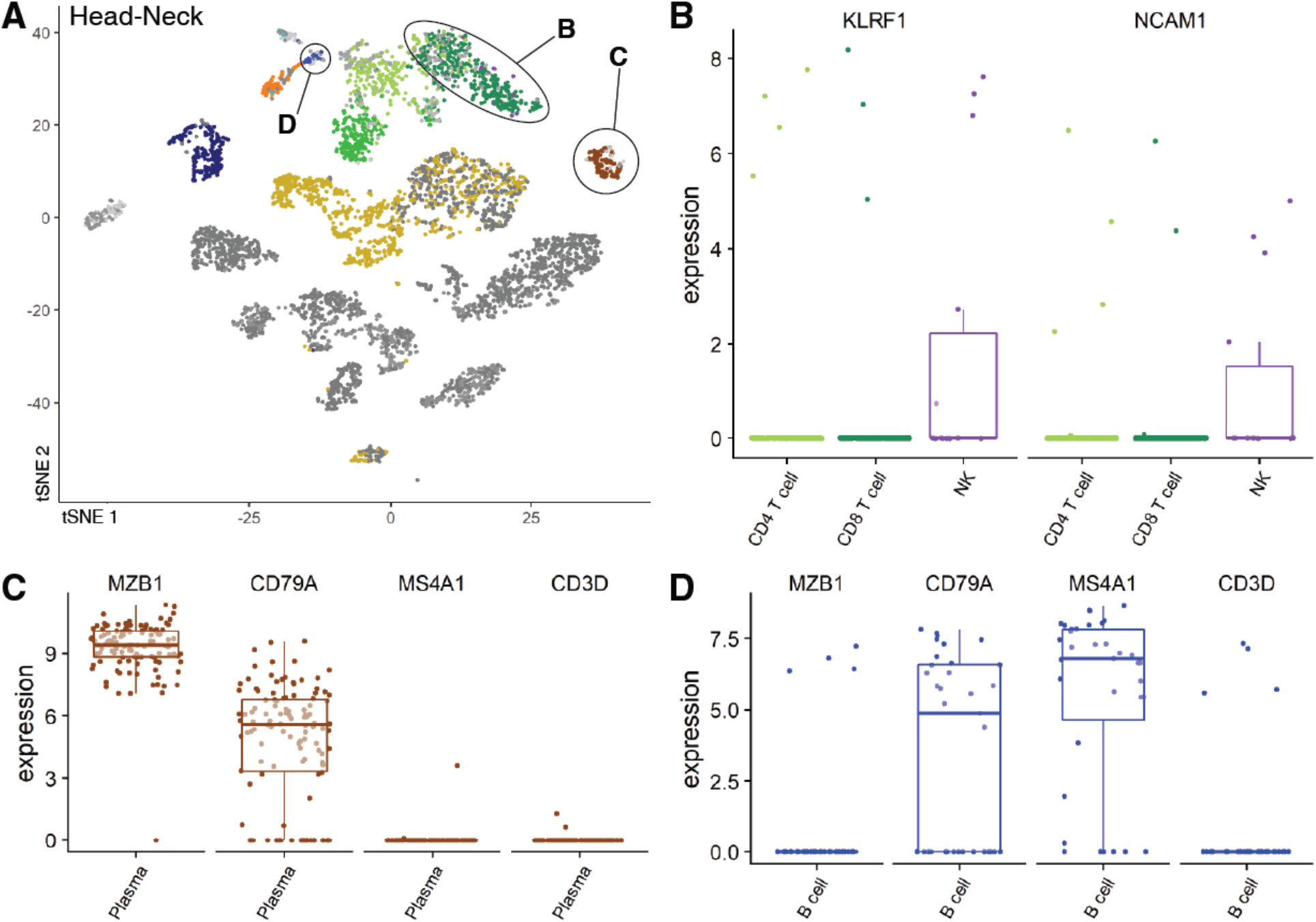
When CHETAH classification differs from the one assigned in the Head-Neck study (Figure 1B, S1), marker gene expression suggests CHETAH’s classification is more plausible. Details as in Figure S3. (**A**) t-SNE plots of the Head-Neck dataset coloured by CHETAH classification. (**B**) Cells previously classified as CD8 T-cells (they cluster together strongly) are probably NK cells. These cells are enriched for NK marker genes *KLRF1* and *NCAM1*. (**C**) Cells previously identified as B-cells are probably *active plasma cells*. Most cells express B-cell marker *CD79A* as well as activation marker *MZB1* while lacking expression of *CD20*/*MS4A1*. (**D**) Cells previously classified as T-cells probably *naive B cells*. Few of these cells express T-cell marker *CD3D*, while most express B cell marker *CD79A*. These cells show no expression of plasma cell marker *MZB1*, but do express *CD20*.

**Figure S5.**
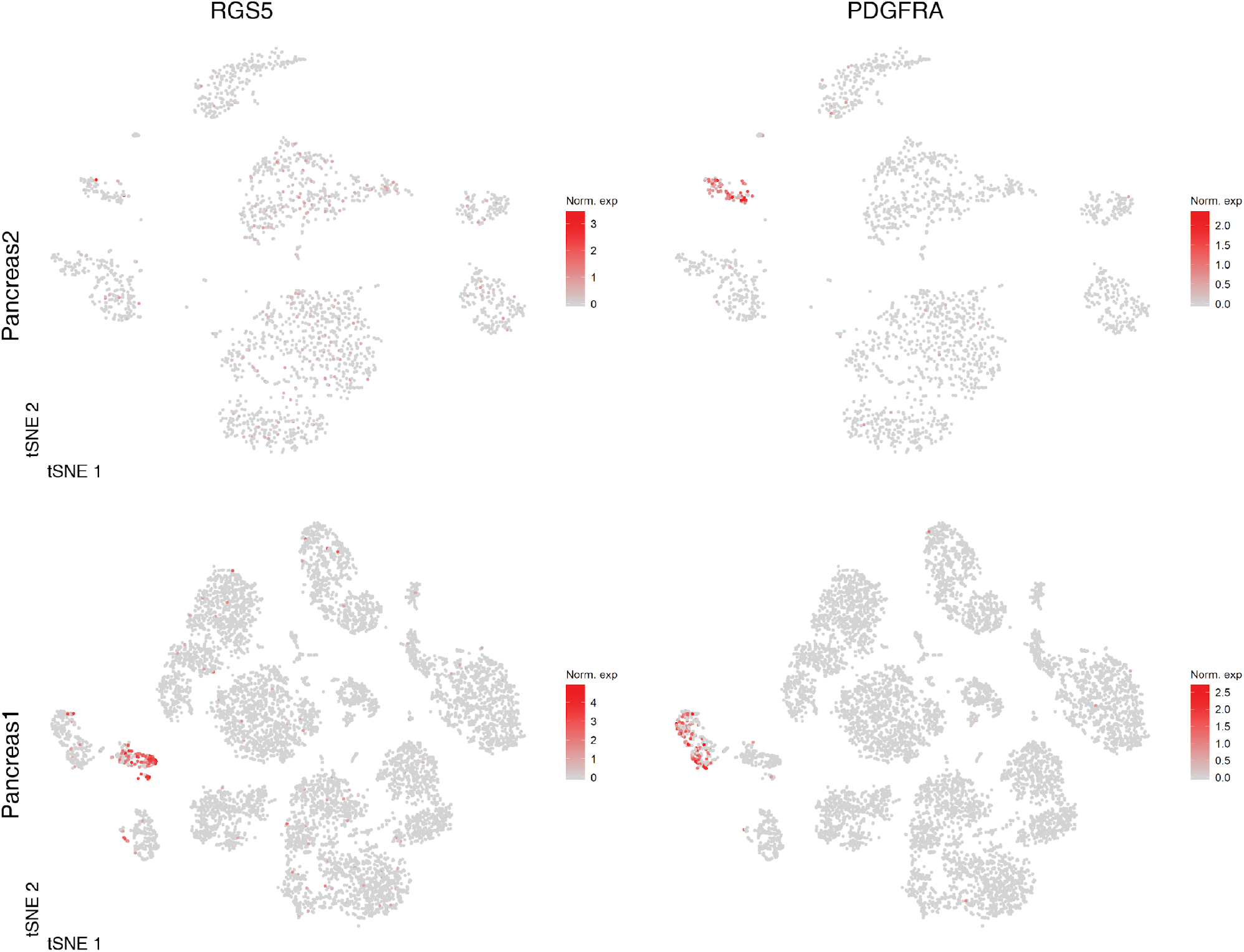
t-SNE plots coloured by the normalized expression of the gene indicated. Rows of the panels: the data classified, as indicated; columns: gene whose expression is shown. In the Pancreas1 publication the *RGS5* gene was used as a marker for *quiescent stellate cells* whereas *PDGFRA* was used as a marker for *activated stellate cells*. The Pancreas2 dataset only has expression of *PDGFRA*, not of *RGS5.*

**Figure S6.**
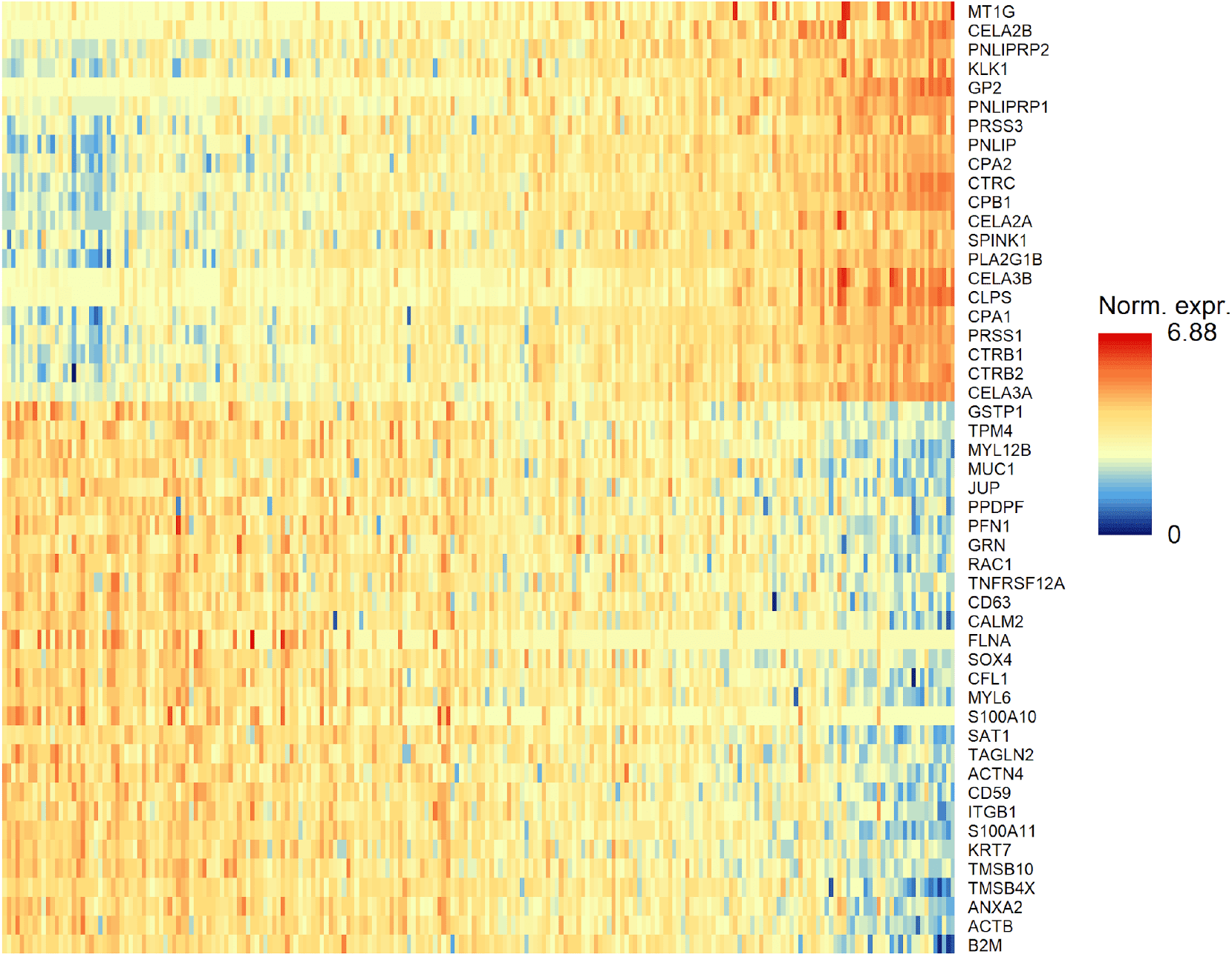
The heatmap shown in main Figure 4D, but with all genes labeled.

**Figure S7.**
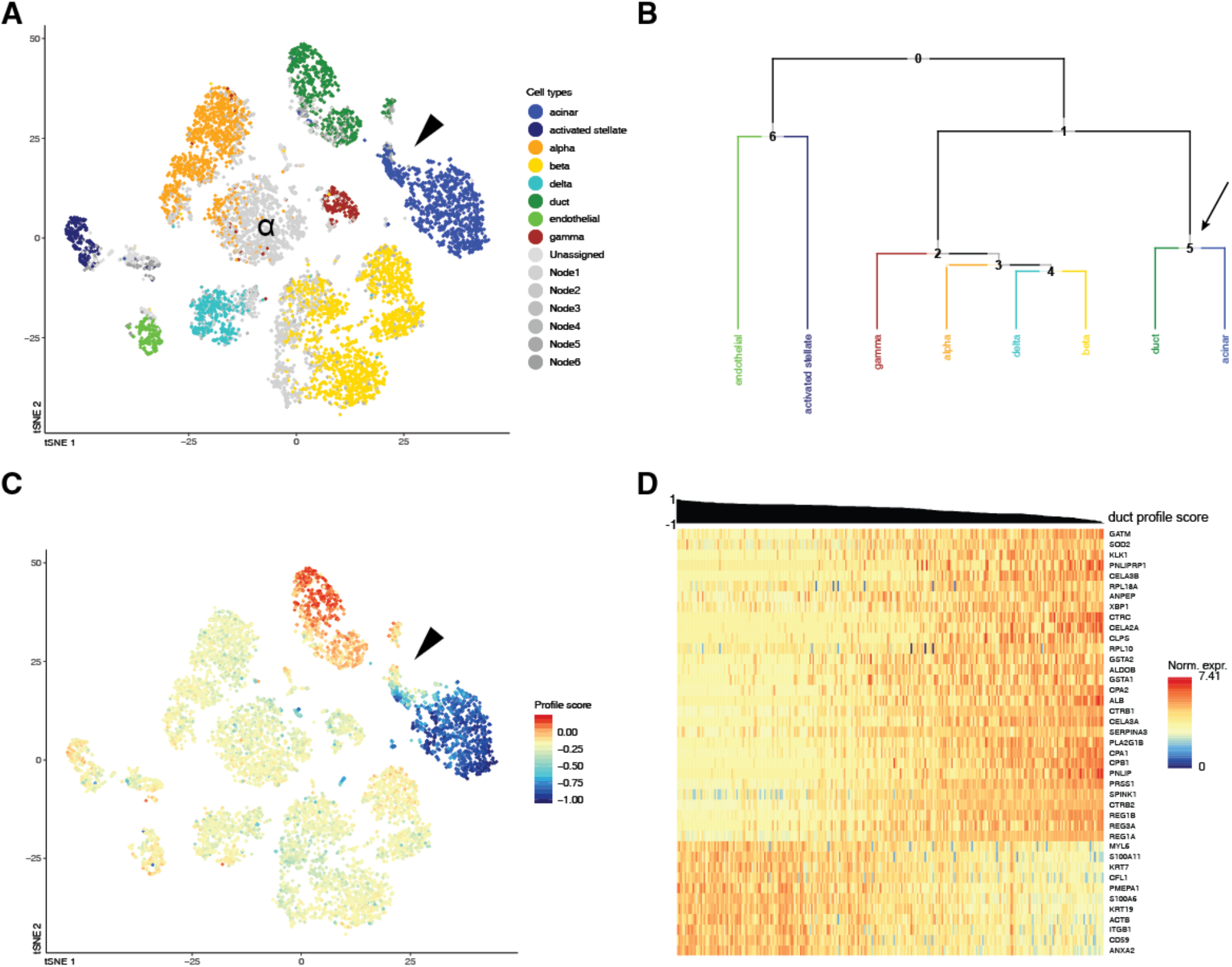
CHETAH identifies opposing gradients of duct and acinar cell marker genes in the Pancreas1 dataset (Baron *et al.*, 2016; reference 10 in main text). (**A**) t-SNE plot of the Pancreas1 dataset as classified by CHETAH, with colours representing the inferred cell types. The arrowpoint indicates a population that was labeled as *duct* cells in the publication, but is classified to a mixture of acinar cells and *intermediate type 6* by CHETAH. (**B**) The classification tree used for A, based on the Pancreas2 dataset. The arrow indicates the *acinar/ductal* intermediate node (Node 5) for which the profile score of duct cells is shown in C. (**C**) As B, but with cells coloured by the profile score for *ductal cell* in Node 5. The cells in the cluster of interest show a gradient of the profile score. (**D**) Heatmap showing the normalized expression of genes used by CHETAH to calculate this profile score. Only the genes that correlate, or anti-correlate more than 0.5 with this profile score in these populations are shown. Rows are genes, columns are cells. The columns are sorted by the duct cell profile score which is shown above the heatmap. For the heatmap with all genes annotated see Figure S6. Note: both CHETAH and scmap have difficulty classifying the alpha cells of one of the donors (as identified in the original publication, here indicated by ‘α’) as they have an unusually low expression of the relevant markers.

